# Interspecific variation in cleaning behaviour and cheating among coral reef cleaner fishes

**DOI:** 10.1101/2025.07.08.663626

**Authors:** Daniele Romeo, Maddalena Ranucci, Melanie Court, Beatriz P. Pereira, José Ricardo Paula, Celia Schunter

## Abstract

Mutualism is an interspecific interaction in which both partners gain a net fitness benefit. However, mutualisms are vulnerable to conflict because one partner may increase its benefits at the other’s expense (cheating). In coral reef cleaning mutualisms, cleaner fishes may cheat by feeding on client mucus rather than ectoparasites, imposing costs on clients and potentially destabilising cooperation. Here, using a standardised experimental framework, we examined cheating behaviour, measured via client jolts as a proxy, in four dedicated (relying on cleaning for sustenance) and three non-dedicated (opportunistic) cleaner fish using an interaction test with three client types (predatory, visitor and resident), and the bystander test, to evaluate potential behavioural changes in the presence of a bystander. When exposed to the same standardised social stimuli, dedicated cleaners responded differently across species, showing pronounced interspecific variation in both interaction time and client jolt frequency, whereas non-dedicated cleaners showed broadly uniform behavioural profiles with rare client jolts. Although captive interactions produced generally lower jolt rates than those typically reported in the wild, cleaner responses still differed across client types, suggesting context-dependent shifts in investment and exploitation with client identity. Bystander effects were weak overall; however, *Labroides bicolor* significantly reduced jolt expression when a bystander was present, suggesting that reputation-related adjustment may be species-specific. These findings highlight the capacity of cleaner species to respond differently to the same social environment, revealing species-specific behavioural strategies with potential consequences for mutualism stability and the selective pressures shaping cooperation–exploitation trade-offs.

## Introduction

Mutualism is an interaction between species in which both partners obtain a net gain in fitness through the reciprocal exchange of resources or services (Bronstein, 1994). This balance, however, can be disrupted when one partner increases its fitness gains at the other’s expense, whose fitness is reduced, thus constituting cheating (Ghoul, Griffin & West, 2014). Cheating can be an important adaptive mechanism for organisms, allowing them to exploit resources and niches otherwise unavailable (Wechsler & Bascompte, 2022). For example, nectar robbers can obtain nectar from flowers whose nectar is normally accessible only to legitimate pollinators (e.g., deep or narrow corollas) by piercing floral tissues, obtaining rewards without providing pollination and imposing costs that can reduce plant fitness (via reward depletion and/or floral damage that lowers visitation) (Irwin, Brody & Waser, 2001; Burkle, Irwin & Newman, 2007). Moreover, cheating can promote phenotypic variation by favouring partner discrimination and control (e.g., partner choice or sanctions), thereby generating coevolutionary feedbacks that diversify interaction strategies and traits, providing a rich substrate for natural selection to act upon (Pellmyr & Leebens Mack, 2000; Wechsler & Bascompte, 2022; Ferrière, Gauduchon & Bronstein, 2007). Despite these insights, it remains unresolved whether cheating follows consistent, species-specific patterns and how these patterns shift across ecological contexts.

In coral reefs, cleaning mutualisms are classic examples of interspecific cooperative interactions, where so-called client fish trade the removal of parasites and dead or infected tissue for an easy meal for cleaner fish (Grutter, 1996). However, in the best-studied system - the Indo-Pacific cleaner wrasses (*Labroides* spp.) - conflict emerges when cleaners consume clients’ mucus instead of parasites, causing tissue damage and imposing an energetic cost because mucus and scales are costly to regenerate (Bshary & Grutter, 2002a). In response to this behaviour, the client imposes partner control mechanisms such as leaving the cleaning station (area of the reef occupied by the cleaner) or displaying aggressive behaviours like chasing the cleaner (Bshary & Grutter, 2002b). This shows that these interactions are not simple by-product mutualisms; instead, clients need to control cleaners’ behaviour to prevent being cheated (i.e., mucus feeding) and to promote cooperative service (i.e., parasite removal) (Bshary & Bronstein, 2011). To mitigate conflict, the cleaner wrasse *Labroides dimidiatus* has developed highly sophisticated social skills, including the use of tactile stimulation for reconciliation and reputation management strategies such as cheating less in the presence of a bystander (Poulin & Grutter, 1996; Pinto et al., 2011). Furthermore, it can adjust quality of service by decreasing cheating while interacting with predators, thereby mitigating predation risk (Bshary, 2002). This high social competence allows *L. dimidiatus* to fine-tune its cheating behaviour to minimise the benefit gained from social interactions (Bshary, 2002; Bshary & Oliveira, 2015). Therefore, cheating in coral reef cleaning behaviour is a driver in the evolution of complex phenotypes, highlighting how conflict can shape sophisticated social strategies and interspecific social behaviours (Ferrière, Gauduchon & Bronstein, 2007; Lin et al., 2024).

The *Labroides* genus comprises five species, all of which are dedicated cleaners that rely on cleaning interactions as their primary trophic strategy throughout life (Vaughan et al., 2017). Despite sharing the same ecological niche, species of this genus exhibit a behavioural continuum of cooperative profiles, ranging from relatively honest cleaning to frequent cheating (Côté & Brandl, 2021; Côté & Mills, 2020; Mills & Côté, 2010). *Labroides rubrolabiatus* and *L. phthirophagus* tend to behave more cooperatively, with minimal partner control (Mills & Côté, 2010; Paula et al pers. communication). *Labroides dimidiatus* adopts an intermediate strategy, employing both cooperative behaviours and manipulative tactics such as tactile stimulation (Bshary & Grutter, 2002b; Grutter, 2004). *Labroides bicolor*, by contrast, cheats frequently, interacts with fewer clients, and for shorter durations (Oates, Manica and Bshary, 2010). Finally, *L. pectoralis* engaged in frequent and long cleaning interactions, suggesting high investment in service provision, while its level of cooperation or cheating remains unquantified (Côté & Brandl, 2021). This interspecific variation may reflect species-specific behavioural tendencies and ecological conditions, such as *L. bicolor*, which shows flexible cooperation levels across its home range, decreasing cheating where repeated interactions with clients are more likely (Oates, Manica and Bshary, 2010). However, although some dedicated cleaners other than *L. dimidiatus* are known to use social tools such as tactile stimulation (Côté & Mills, 2020), their capacity to adjust service quality, and thereby reduce or increase cheating, across social contexts (e.g., client identity and the presence or absence of a bystander) remains poorly understood. This makes it unclear whether reputation-based or context-dependent strategies are widespread within *Labroides* or largely species-specific, a question that remains largely untested in controlled comparative frameworks exposing multiple cleaner species to the same social stimuli. Even less is known about non-dedicated cleaners, which engage in cleaning only during certain life stages or as a supplementary feeding strategy (Vaughan et al., 2017). Since these species rely on alternative food sources, selection for cleaning-related behavioural specialisation may be weaker, potentially leading to lower service investment and a reduced ability to fine-tune service quality across social contexts (e.g., weaker context-dependent modulation of cheating with client identity or bystander presence) (Barbu et al., 2011; Bshary & Oliveira, 2015). Overall, these gaps make a controlled interspecific study necessary not only to test whether context-dependent service modulation is general or species-specific, but also to compare how behavioural responses to the same social stimuli vary across cleaner species with different ecological dependence on cleaning.

Here, we investigate the interspecific variation in cheating occurrence and frequency, measured via client jolts as a behavioural proxy, among dedicated and non-dedicated cleaner wrasses using two behavioural tests: an interaction test, which compares interactions with three different client types (predators, residents, and non-residents), and a bystander test, which examines interactions in the presence and absence of a bystander. We predict that variation in cheating will be more pronounced among dedicated cleaners, reflecting different cleaning strategies adopted by each species, while non-dedicated cleaners are expected to show less variability due to their more opportunistic use of cleaning. By focusing on these behavioural patterns, this study aims to advance our understanding of the ecological and evolutionary dynamics that shape mutualistic interactions in coral reef ecosystems and drive behavioural diversity and species adaptation.

## Material and Methods

To investigate interspecific variation in cheating among cleaner wrasse species, four species of dedicated cleaners and three species of non-dedicated cleaners were tested via an interaction test with three different clients and a bystander test (Pinto et al., 2011) (Table 1, Suppl. Table 1). Dedicated cleaner species were selected from the genus *Labroides*, as it is the most studied group of dedicated cleaners (Côté & Brandl, 2021; Côté & Mills, 2020). Non-dedicated cleaner species were chosen based on their phylogenetic proximity to *Labroides* and previous knowledge on performing cleaning behaviours (Vaughan et al. 2017; Sazima, Moura & Gasparini, 1998; Baliga & Law, 2016; Baliga & Mehta, 2019). The behavioural tests were conducted at Laboratório Marítimo da Guia in Cascais (Portugal) and The Swire Institute of Marine Science (SWIMS) in Hong Kong. Within this study we used *Dascyllus trimaculatus*, *Zebrasoma scopas*, and *Paracirrhites forsteri* as client fish. The fishes were provided by the local suppliers Tropical Marine Centre (TMC; Portugal) and by Sealife Hong Kong limited (Hong Kong). Fish were deparasitised by the commercial suppliers prior to delivery, then acclimated for three days under laboratory conditions and fed ad libitum with mysids (*Mysis* spp.) and copepods twice daily throughout acclimation. In total, we tested 65 individuals; species-specific sample sizes are reported in Table 1.

**Table 1.**
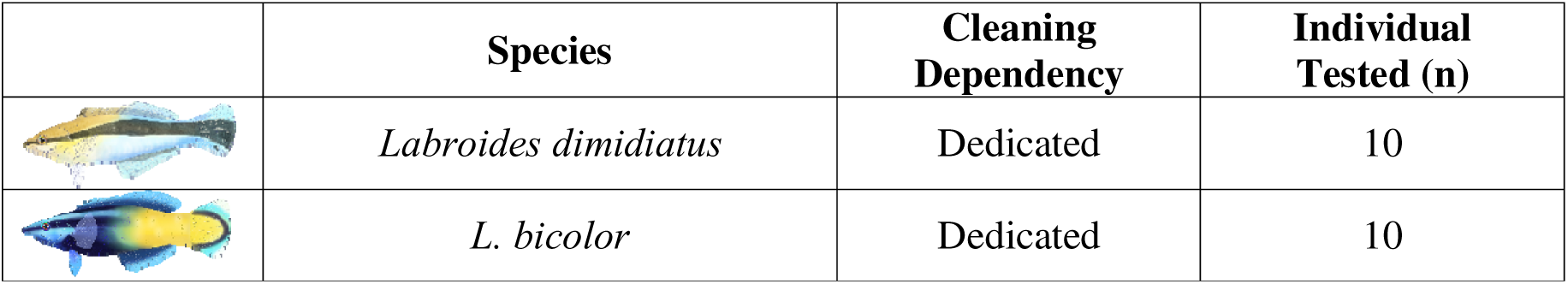
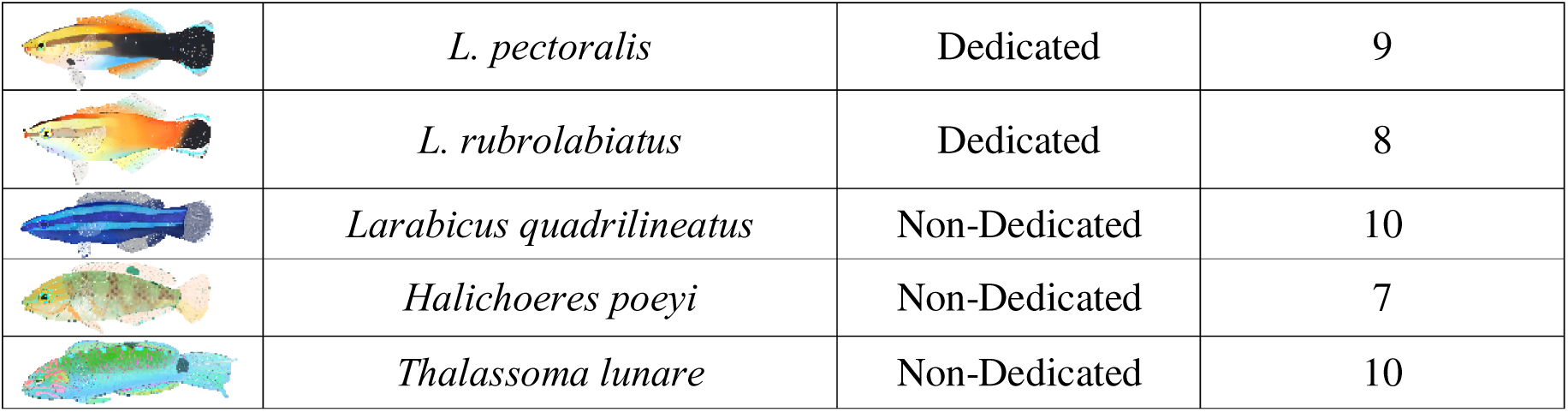
Cleaner wrasse species included in the study: four dedicated Labroides species and three non-dedicated cleaners.

|  | Species | Cleaning Dependency | Individual Tested (n) |
| --- | --- | --- | --- |
|  | *Labroides dimidiatus* | Dedicated | 10 |
|  | *L. bicolor* | Dedicated | 10 |
|  | *L. pectoralis* | Dedicated | 9 |
|  | *L. rubrolabiatus* | Dedicated | 8 |
|  | *Larabicus quadrilineatus* | Non-Dedicated | 10 |
|  | *Halichoeres poeyi* | Non-Dedicated | 7 |
|  | *Thalassoma lunare* | Non-Dedicated | 10 |

### Interaction Test

On the fourth day after arrival to the laboratory, an interaction test was carried out where each cleaner interacted with the client in the same tank. Food was withheld on the morning of the experiment day to maintain motivation for interactions, following standard practice in similar assays (de Abreu et al., 2018; Paula et al., 2019; Ramírez-Calero et al., 2022; 2023). To evaluate the variation in cheating in the study species, three different types of clients were selected: a resident (*Dascyllus trimaculatus*), a non-resident (*Zebrasoma scopas*) and a predator client (*Paracirrhites forsteri*). The client species were chosen due to their widespread spatial overlap with all the cleaner species in the study. On day four, the cleaner and the client were placed on opposite sides of an experimental tank (50 x 31 x 40 cm), separated by two partitions: one opaque to avoid visual contact and prevent the cleaner from becoming stressed by seeing the client (especially the predator), and one transparent partition (Fig. 1A). After 30 minutes of acclimation to the experimental tank, the opaque partition was removed to allow five minutes of visual acclimation. Afterwards, the transparent partition was removed, and the interaction was recorded using a GoPro Hero Black 10 for 45 minutes. Then, the partitions were placed back in the experimental tank, the client was removed, and a new client was introduced into the experimental tank for another 30 minutes of acclimation. To ensure comparability across individuals and minimise uncontrolled variation due to order effects, the sequence of client presentation was kept constant: predator, non-resident, and resident. The predator was presented first to reduce the risk of heightened stress responses that might occur if it were introduced after other interactions. The resident client was always presented last, because it is a common, predictable client in the wild and helped reduce stress at the end of the trial, allowing the cleaner to finish the session in a more relaxed state.

**Figure 1.**
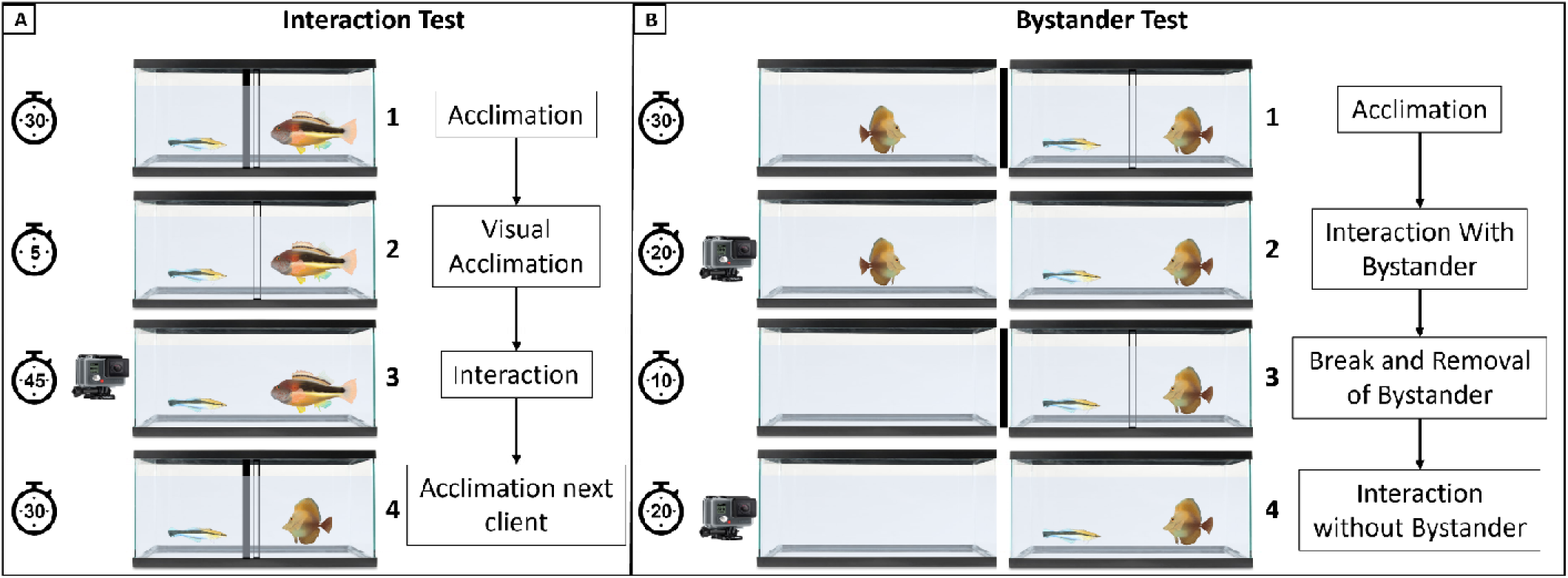
Experimental design for the Interaction Test (A) and the Bystander Test (B). For the Interaction Test (A), the cleaner and client fish are placed together in the experimental tank, separated by an opaque and transparent barrier (1. A). After 30 minutes, the opaque barrier is removed (2. A), followed by the transparent barrier after 5 minutes (3. A), allowing interaction. Their behaviour is recorded for 45 minutes. The client is then replaced, and the process is repeated with the next client (4. A). For the Bystander Test (B), the cleaner and client are initially separated by a transparent barrier in the experimental tank, with a bystander fish in an adjacent tank separated by an opaque barrier (1. B). After 30 minutes, all barriers are removed, and interactions are recorded for 20 minutes (2. B). The barriers are then put back in place, the bystander is removed (3. B), and once the barriers are removed again, interactions are recorded for another 20 minutes (4. B).

### Bystander Test

On day five, the fish were tested using the bystander test, following a protocol adapted from Pinto et al., 2011 (Pinto et al., 2011). The cleaner was placed in the experimental tank with the client *Zebrasoma scopas*, separated by a transparent partition (Fig. 1B). Another individual of *Z. scopas* was placed in an adjacent tank, separated by an opaque partition. After 30 minutes of acclimation, all partitions were removed, allowing the fish in the same tank to interact and the bystander to be visible. The interaction was recorded for 20 minutes to assess the cleaner’s behaviour in the presence of the bystander. Then, the partitions were placed back for 10 minutes, and the bystander fish was removed. Finally, the partitions were removed again, and the interaction, now without the bystander, was recorded for another 20 minutes to compare the cleaner’s behaviour in the absence of a bystander.

### Supervised machine learning video analysis

All the videos produced were processed to have the same resolution and length (1920×1080; 30 fps; 40 minutes), with the first 5 minutes removed to exclude effects from partition removal and other minor set-up adjustments at trial start, ensuring standardised video length and resolution across recordings. SimBA (v.2.3.5) (Goodwin et al., 2024), an open-source package that uses pose-estimation to create supervised machine learning classifiers of social behaviour, was used to analyse variations in cheating and time spent interacting by the fish in the two behavioural tests. Pose-estimation data were generated using DeepLabCut (v.2.3.9) (Nath et al., 2019), creating 21 models (one for each species tested per client). Approximately 400 frames were labelled for each model to ensure a good representation of behavioural variation across videos. A ResNet-50-based neural network was trained with default parameters for 500k iterations (batch size 1), chosen for its optimal balance of high accuracy in distinguishing complex poses and computational efficiency, making it particularly effective for multi-animal tracking in scenarios with overlapping or interacting subjects. The models were validated using two shuffles; mean training and test errors were 3.68 and 4.94 pixels, respectively (range across species-client models: training 3.42–3.97; test 4.80–5.11 pixels). DeepLabCut models return X–Y coordinates and a likelihood score (0–1) for each detection. A p-cutoff of 0.6 was applied to exclude low-confidence detections and retain only coordinates with likelihood ≥0.6 for downstream analyses, ensuring sufficient data quality while excluding unreliable predictions, aligned with standard practices in pose-estimation tasks. The behavioural annotation required for training the machine learning model in SimBA was produced using the software BORIS (v.7.13.8) (Friard & Gamba, 2016), focusing on two key behaviours: client jolts (rapid movements of the client away from the species after receiving a bite) and interaction time (defined as the duration in which the cleaner fish was actively inspecting or cleaning the client). Client jolts were used as a behavioural proxy for cheating, following established conventions in cleaner fish research (e.g., Bshary and Grutter, 2002; Pinto et al., 2011). The pose-estimation data from DeepLabCut and the behavioural annotations from BORIS were then used to train supervised machine learning models in SimBA using the Random Forest algorithm (2000 estimators; split criterion: entropy; max features: sqrt; train–test split: 0.2). Following model training (F1-scores: 0.80 for interaction time and 0.52 for client jolts, Supp. Fig. 1), validation videos were generated and discrimination thresholds (i.e., the minimum prediction score produced by the model above which a frame is classified as the behaviour) were set to 0.45 for interaction time and 0.73 for client jolts. For the client jolts classifier, performance showed very high recall (0.998) paired with low precision (0.352), indicating reliable detection of true positives while generating many false positives. Therefore, short clips were generated for each predicted jolt event and all false positives were manually excluded from the final dataset.

### Statistical Analysis

For the interaction test, data obtained from supervised machine learning were analysed in R (v4.3.1) (R Core Team, 2023). Because the variable client jolts was highly zero-inflated, and some species never produced any jolts with specific clients, causing complete separation (i.e., zero counts for all observations in some species–client combinations) and unreliable parameter estimates, a two-part (hurdle-like (Cragg, 1971; Albert & Anderson, 1984) modelling strategy was adopted. First, to test whether the probability of observing at least one client jolt differed among cleaner species and between client types, both overall and within each client, client-jolt occurrence was modelled as a binary response (AnyJolt; 1 = at least one client jolt observed within a video; 0 = no jolts) using a bias-reduced binomial logistic regression in brglm2 (Kosmidis, 2021), with Species, Client, and their interaction as predictors (AnyJolt ∼ Species * Client) (Table 2). Second, to quantify variation in jolt rates among species and client types, jolt counts were analysed only for videos with at least one jolt using a negative binomial GLM (MASS; Ripley et al., 2013), with total interaction time per video included as an offset to express rates per 100 s of interaction (Cheating ∼ Species * Client + offset(log(Interaction/100))) (Table 2). Interaction time was analysed using a Gamma model with log link implemented in glmmTMB (Brooks et al., 2017) (Interaction ∼ Species * Client) (Table 2). For each model, Species, Client, and Species × Client effects were assessed using likelihood-ratio χ² tests by comparing the full model with reduced models obtained by removing one term at a time (drop1) (Supp. Table 2). Pairwise comparisons among species within each client were performed using emmeans (Lenth, 2024), with p-values adjusted using the Benjamini–Hochberg procedure, and model diagnostics were evaluated using DHARMa (Hartig, 2024) (Supp. Fig. 2). Differences in variance between dedicated and non-dedicated cleaners were assessed using Levene’s test (center = median) (Levene, 1960), as implemented in the R package car (Fox & Weisberg, 2018), for total interaction time and client jolts per 100 s of interaction. For the bystander test, within each species, behavioural differences between the presence and absence of the bystander were assessed using two-tailed Wilcoxon signed-rank tests (Wilcoxon, 1945) implemented in base R (stats::wilcox.test; R Core Team, 2023) for the same variables analysed in the interaction test, client jolts per 100 s of interaction and interaction time.

**Table 2.** The table reports fixed-effect parameter estimates (Estimate), standard errors (SE), Wald z-statistics (z value), and associated two-sided p-values (Pr(>|z|)) for models used in this study. Estimates are reported on the link scale (log-odds for binomial models; log scale for negative binomial and Gamma models). For categorical predictors, coefficients are expressed relative to the reference levels; therefore, the intercept corresponds to the expected response for the reference Species–Client combination. Reference levels were H. poeyi and Non Resident for the binomial and Gamma models, whereas for the negative binomial model (Cheating > 0) the intercept corresponds to L. bicolor and Non Resident. Significant p-values (Pr(>|z|) < 0.05) are shown in bold.

| **Model:** AnyJolt ~ Client * Species (brglm2, bias-reduced binomial logit; AS_mean). |  |  |  |  |
| --- | --- | --- | --- | --- |
| **Response:** AnyJolt (0/1 per video; $\geq 1$ jolt vs none) | | | | |
| Term | Estimate | Std. Error | z value | Pr(>\|z\|) |
| (Intercept) | -2.56495 | 1.585188 | -1.61807 | 0.105647 |
| ClientPredator | -0.1431 | 2.225065 | -0.06431 | 0.948721 |
| ClientResident | -0.1431 | 2.225065 | -0.06431 | 0.948721 |
| SpeciesL. bicolor | 3.32709 | 1.724463 | 1.929348 | 0.053688 |
| SpeciesL. dimidiatus | 1.80281 | 1.724463 | 1.045432 | 0.295823 |
| SpeciesL. pectoralis | 2.364279 | 1.720975 | 1.373802 | 0.169503 |
| SpeciesL. quadrilineatus | -0.37949 | 2.202726 | -0.17228 | 0.863216 |
| SpeciesL. rubrolabiatus | 2.112965 | 1.743214 | 1.212108 | 0.225471 |
| SpeciesT. lunare | 0.830349 | 1.839641 | 0.451364 | 0.651727 |
| ClientPredator:SpeciesL. bicolor | 2.425483 | 2.777882 | 0.873141 | 0.382586 |
| ClientResident:SpeciesL. bicolor | -0.98676 | 2.413616 | -0.40883 | 0.682663 |
| ClientPredator:SpeciesL. dimidiatus | 1.105911 | 2.420909 | 0.456816 | 0.647803 |
| ClientResident:SpeciesL. dimidiatus | -2.0392 | 2.784071 | -0.73245 | 0.463893 |
| ClientPredator:SpeciesL. pectoralis | 0.343771 | 2.428961 | 0.14153 | 0.887451 |
| ClientResident:SpeciesL. pectoralis | -2.60067 | 2.781911 | -0.93485 | 0.349866 |
| ClientPredator:SpeciesL. quadrilineatus | 1.241713 | 2.852932 | 0.435241 | 0.663388 |
| ClientResident:SpeciesL. quadrilineatus | 1.352939 | 2.856844 | 0.473578 | 0.635801 |
| ClientPredator:SpeciesL. rubrolabiatus | -0.87125 | 2.53271 | -0.344 | 0.730847 |
| ClientResident:SpeciesL. rubrolabiatus | -2.11296 | 2.813358 | -0.75105 | 0.452624 |
| ClientPredator:SpeciesT. lunare | -0.95551 | 2.86439 | -0.33358 | 0.738694 |
| ClientResident:SpeciesT. lunare | -1.06674 | 2.856844 | -0.3734 | 0.708853 |
| **Model:** Cheating ~ Client * Species + offset(log(Interaction/100)) (negative binomial GLM; MASS; subset Cheating > 0) |  |  |  |  |
| **Response:** Cheating (count of client jolts per video; analysed only when Cheating > 0; offset expresses jolts per 100 s). |  |  |  |  |
| Term | Estimate | Std. Error | z value | Pr(>\|z\|) |
| (Intercept) | 0.198868 | 0.244627 | 0.812944 | 0.41625 |
| ClientPredator | 0.499979 | 0.334629 | 1.494132 | 0.135141 |
| ClientResident | -1.60625 | 0.522978 | -3.07135 | **0.002131** |
| SpeciesL. dimidiatus | -1.23203 | 0.71507 | -1.72295 | 0.084898 |
| SpeciesL. pectoralis | -1.78176 | 0.54791 | -3.25193 | **0.001146** |
| SpeciesL. quadrilineatus | 0.502756 | 1.033496 | 0.486461 | 0.62664 |
| SpeciesL. rubrolabiatus | -0.81697 | 0.498511 | -1.63881 | 0.101253 |
| SpeciesT. lunare | -0.94223 | 1.18924 | -0.7923 | 0.428187 |
| ClientPredator:SpeciesL. dimidiatus | 0.621585 | 0.840704 | 0.739362 | 0.459687 |
| ClientPredator:SpeciesL. pectoralis | 1.389107 | 0.76367 | 1.818988 | 0.068913 |
| ClientPredator:SpeciesL. quadrilineatus | -1.49838 | 1.573117 | -0.95249 | 0.340847 |
| ClientPredator:SpeciesL. rubrolabiatus | -0.89697 | 0.924715 | -0.96999 | 0.33205 |
| **Model:** Interaction ~ Client * Species (Gamma model, log link; glmmTMB) |  |  |  |  |
| **Response:** Interaction (total interaction time per video; seconds). |  |  |  |  |
| Term | Estimate | Std. Error | z value | Pr(>\|z\|) |
| (Intercept) | 4.744687 | 0.382632 | 12.40012 | **2.61E-35** |
| ClientPredator | -0.37917 | 0.52144 | -0.72717 | 0.467125 |
| ClientResident | -0.25152 | 0.521439 | -0.48236 | 0.629548 |
| SpeciesL. bicolor | 2.492807 | 0.483996 | 5.150467 | **2.6E-07** |
| SpeciesL. dimidiatus | 1.029275 | 0.483997 | 2.126617 | **0.033452** |
| SpeciesL. pectoralis | 1.875501 | 0.493977 | 3.796735 | **0.000147** |
| SpeciesL. quadrilineatus | 0.327548 | 0.493977 | 0.663082 | 0.507278 |
| SpeciesL. rubrolabiatus | 1.702195 | 0.506176 | 3.362851 | **0.000771** |
| SpeciesT. lunare | 0.370358 | 0.493977 | 0.749746 | 0.453407 |
| ClientPredator:SpeciesL. bicolor | -0.6019 | 0.669021 | -0.89967 | 0.368298 |
| ClientResident:SpeciesL. bicolor | -0.65077 | 0.669021 | -0.97271 | 0.330696 |
| ClientPredator:SpeciesL. dimidiatus | 0.023253 | 0.676277 | 0.034384 | 0.972571 |
| ClientResident:SpeciesL. dimidiatus | -0.1272 | 0.676276 | -0.18808 | 0.85081 |
| ClientPredator:SpeciesL. pectoralis | -1.63093 | 0.692324 | -2.35574 | **0.018486** |
| ClientResident:SpeciesL. pectoralis | -2.42135 | 0.683455 | -3.54281 | **0.000396** |
| ClientPredator:SpeciesL. quadrilineatus | 0.254768 | 0.676277 | 0.376721 | 0.706381 |
| ClientResident:SpeciesL. quadrilineatus | 0.38032 | 0.683455 | 0.556467 | 0.577891 |
| ClientPredator:SpeciesL. rubrolabiatus | -0.52441 | 0.712179 | -0.73635 | 0.46152 |
| ClientResident:SpeciesL. rubrolabiatus | -2.24374 | 0.71218 | -3.15053 | **0.00163** |
| ClientPredator:SpeciesT. lunare | -0.4802 | 0.692323 | -0.6936 | 0.487931 |
| ClientResident:SpeciesT. lunare | -0.12443 | 0.683456 | -0.18205 | 0.85554 |

## Results

### Interaction test

The interaction test revealed significant interspecific differences in both cleaning and cheating behaviours, with dedicated cleaners showing significantly greater behavioural variability (from 0 to 8.60 jolts per 100 s; interaction times ranging from 0 to 2358 s) compared to non-dedicated cleaners (from 0 to 0.74 jolts per 100 s; interaction times ranging from 0 to 546 s) (Levene’s test: interaction time, F = 27.47, p = 4.47 × 10 ; jolts per 100 s, F = 11.41, p = 8.98 × 10 ; Fig. 2). Consistently, client-jolt occurrence and, when present, jolt rates were significantly affected by both species and client type; interaction time was also significantly affected by species and client type and showed a significant client-by-species interaction (drop-test LRT: AnyJolt—Client χ²=21.32, p=2.35×10 ; Species χ²=59.36, p=6.09×10 ¹¹; Cheating|>0—Client χ²=18.46, p=9.81×10 ; Species χ²=11.73, p=0.039; Interaction time—Client χ²=28.30, p=7.17×10 ; Species χ²=67.48, p=1.34×10 ¹²; Client×Species χ²=34.88, p=4.89×10 ; Supp. Table 2).

**Figure 2.**
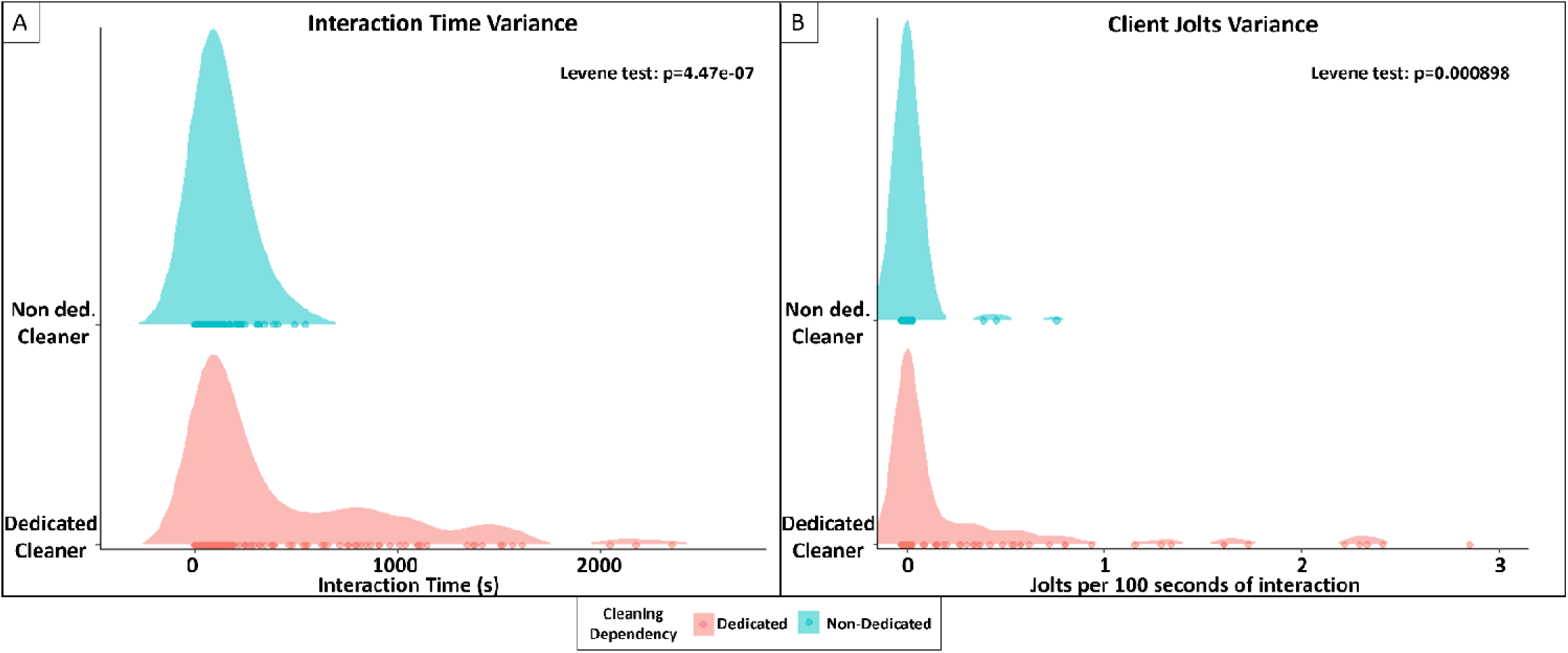
Differences in behavioural variability between dedicated and non-dedicated cleaners. Kernel density plots show the distribution of (A) interaction time and (B) client jolts per 100 s of interaction for dedicated (pink) and non-dedicated (blue) cleaners; points along the baseline represent individual observations. Differences in variance between groups were assessed using Levene’s test (center = median), showing significantly greater variability in dedicated cleaners for both interaction time (p = 4.47 × 10□□) and jolts per 100 s (p = 8.98 × 10□□).

At the species level, *Labroides bicolor* was associated with the highest client jolt occurrence (≥1 jolt per video), with at least one client jolt observed in 70% of videos, which differed significantly from all other species (all p ≤ 0.03; Fig. 3, Supp. Table 3). Furthermore, client jolts were mostly found with dedicated cleaners and differed in their distribution across client types. Predator clients exhibited jolts in all *L. bicolor* videos (100%), at a mean rate of 2.07 ± 1.14 jolts per 100 s. Jolts were also observed in *L. dimidiatus* (55.6% of predator videos; 0.81 ± 1.78) and *L. pectoralis* (44.4%; 1.24 ± 2.80) videos, however were rare in *L. rubrolabiatus* (14.3%; 0.05 ± 0.14) (Fig. 4, Supp. Table 3, 4). Non-resident clients showed a similar ranking, jolting most frequently when cleaned by *L. bicolor* (70% of videos; 0.85 ± 0.92) and less frequently when cleaned by *L. dimidiatus* (30%; 0.11 ± 0.19), *L. pectoralis* (44.4%; 0.13 ± 0.22), and *L. rubrolabiatus* (37.5%; 0.19 ± 0.40) (Fig. 4, Supp. Table 3, 4). Resident clients were less likely to jolt (8.3% of videos) than non-resident (29.5%) and predator clients (35.6%), although these differences were not statistically significant (non-resident vs resident: p = 0.0956; predator vs resident: p = 0.0756), with jolts occurring only with *L. bicolor* (40% of videos; 0.10 ± 0.14) and *L. quadrilineatus* (10%; 0.04 ± 0.13) (Fig. 4, Supp. Table 3, 4). Among non-dedicated cleaners, jolts were observed only sporadically, occurring in two videos in *L. quadrilineatus* (one with a predator client and one with a resident client) and in one video in *T. lunare* (non-resident client), and were absent in *H. poeyi* (Fig. 4, Supp. Table 3, 4). However, after post hoc correction, only *L. bicolor* vs *L. pectoralis* (non-resident clients) remained significant (rate ratio = 5.94, p = 0.011), whereas all other contrasts were non-significant (p ≥ 0.278) (Supp. Table 3, 4).

**Figure 3.**
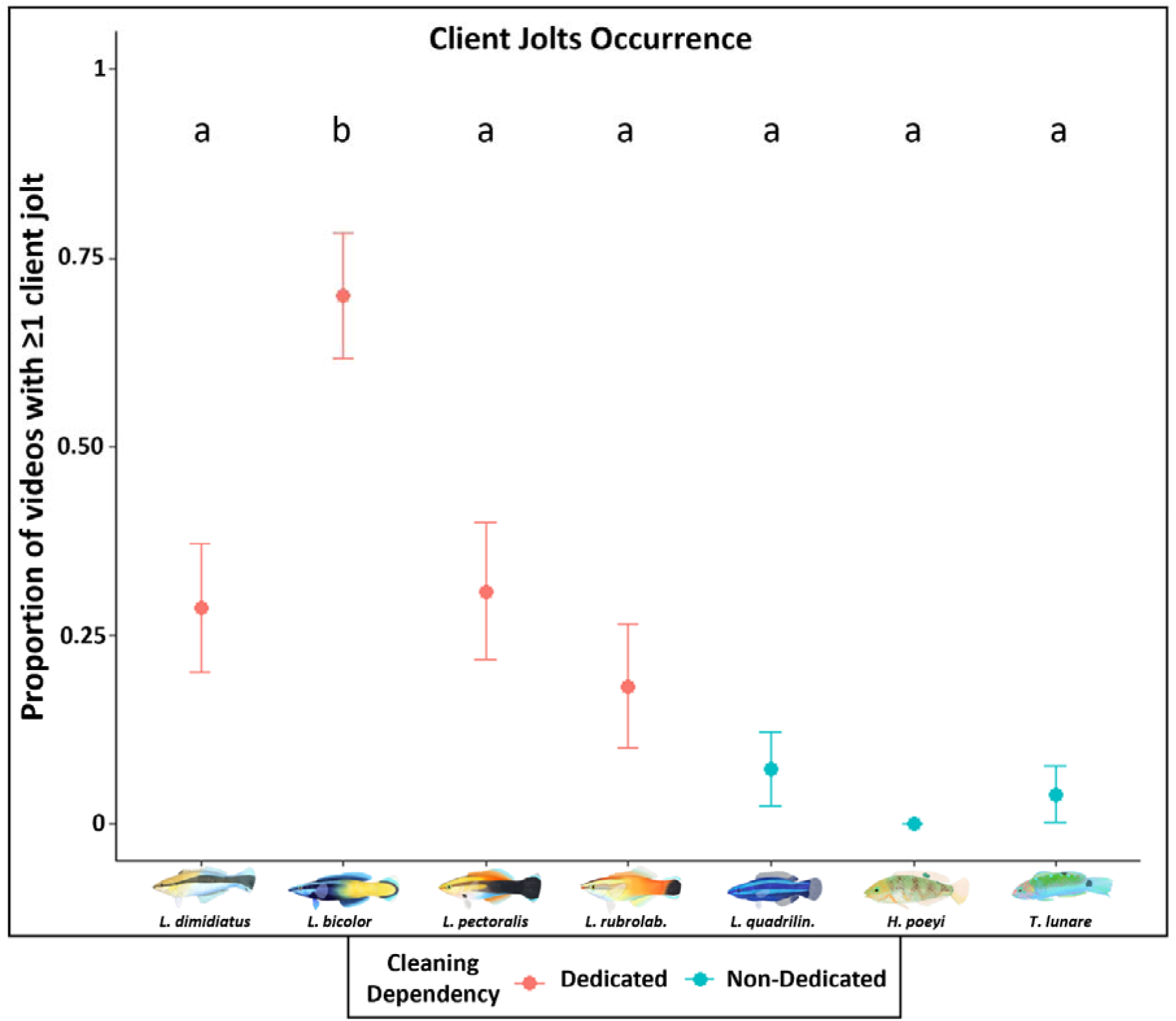
Variation in client jolt occurrence across cleaner fish species. Points show the proportion of videos with ≥1 client jolt, with error bars indicating binomial SE. Colours denote cleaning dependency (Dedicated vs Non-dedicated). Different lowercase letters indicate significant differences among species, based on post hoc comparisons (emmeans, BH-corrected) from a bias-reduced binomial logistic regression (brglm2).

**Figure 4.**
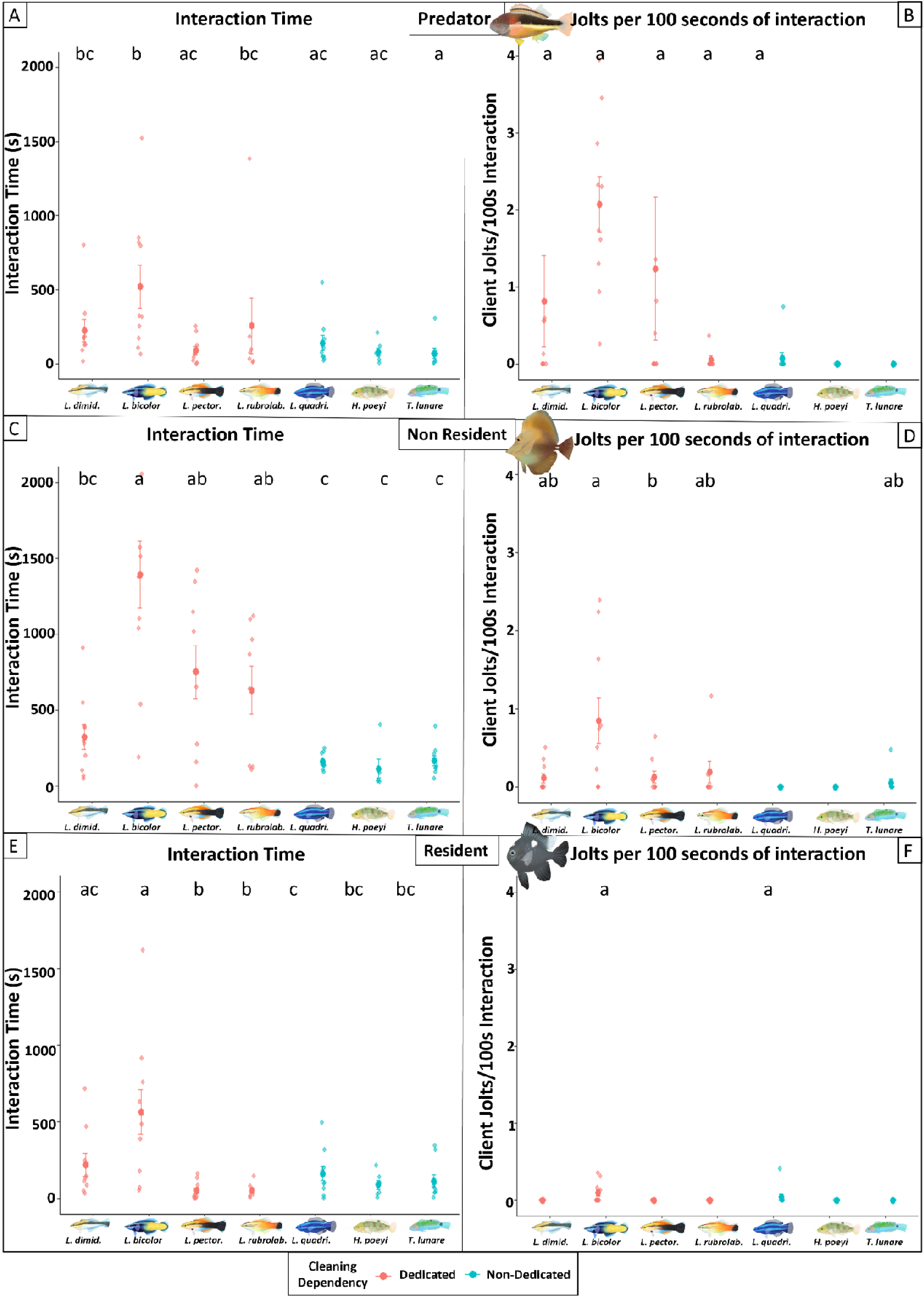
Variation in interaction time and client jolts per 100 s of interaction across cleaner fish species and client types. Panels show (A, C, E) interaction time (s) and (B, D, F) client jolts per 100 s of interaction for seven cleaner fish species interacting with three client types: predators (A–B), non-residents (C–D), and residents (E–F). Small points represent individual observations, while larger points and error bars indicate the mean ± SE for each species. Colours denote cleaning dependency (Dedicated vs Non-dedicated). Different lowercase letters indicate significant differences among species within each panel, based on post hoc comparisons (emmeans, BH-corrected). Interaction time was analysed with a Gamma model, whereas jolt rates were analysed with a negative binomial model fitted only to videos in which at least one jolt occurred.

Interaction times also differed across species and client types. Dedicated cleaners spent the longest time interacting with non-resident clients (mean: 781.64 s; range = 0.97–2358 s), compared with predator (281.00 s; 0–1519.27 s) and resident clients (241.55 s; 0.77–1615.43 s), whereas non-dedicated cleaners showed shorter and more uniform interaction times across clients (151.01 s; 27.63–404.27 s with non-residents; 100.94 s; 2.57–546.17 s with predators; and 126.46 s; 0–494.20 s with residents) (Fig. 4). Across all client types, *L. bicolor* showed the longest interactions (non-resident: 1390.61 ± 700.69 s; predator: 521.35 ± 462.23 s; resident: 564.08 ± 466.76 s; Fig. 4), differing significantly from non-dedicated cleaners across all client types (*H. poeyi*: non-resident p = 5.20 × 10□□, predator p = 4.45 × 10□□, resident p = 4.66 × 10□□; *L. quadrilineatus*: non-resident p = 5.20 × 10□□, predator p = 0.00943, resident p = 0.0197; *T. lunare*: non-resident p = 5.80 × 10□□, predator p = 1.42 × 10□□, resident p = 0.00110) (Fig. 4, Supp. Table 5). It also differed from *L. dimidiatus* with non-resident clients (321.81 ± 260.50 s; p = 0.00168) and from *L. pectoralis* and *L. rubrolabiatus* with resident clients (51.80 ± 59.93 s, p = 6.17 × 10□□; 52.02 ± 48.07 s, p = 2.59 × 10□□) (Fig. 4, Supp. Table 5). Furthermore, consistent with greater variability among dedicated cleaners, L. dimidiatus interacted longer with resident clients (220.36 ± 227.53 s) than *L. pectoralis* and *L. rubrolabiatus* (p = 0.00440 and p = 0.00784) (Fig. 4, Supp. Table 5). Finally, no significant differences were detected in pairwise comparisons among non-dedicated cleaners (Fig. 4, Supp. Table 5).

### Bystander test

Bystander effects were weak across species and behavioural variables (Fig. 3, Supp. Table 6). Only *L. bicolor* elicited significantly fewer client jolts when a bystander was present than when it was absent (p = 0.028; Fig. 5, Supp. Table 6). *L. dimidiatus* also tended to provoke fewer jolts with a bystander present; however, this effect was not significant (p = 0.249). Interaction time remained similar between conditions, with no significant differences detected in any species (Fig. 5A, Supp. Table 6).

**Figure 5:**
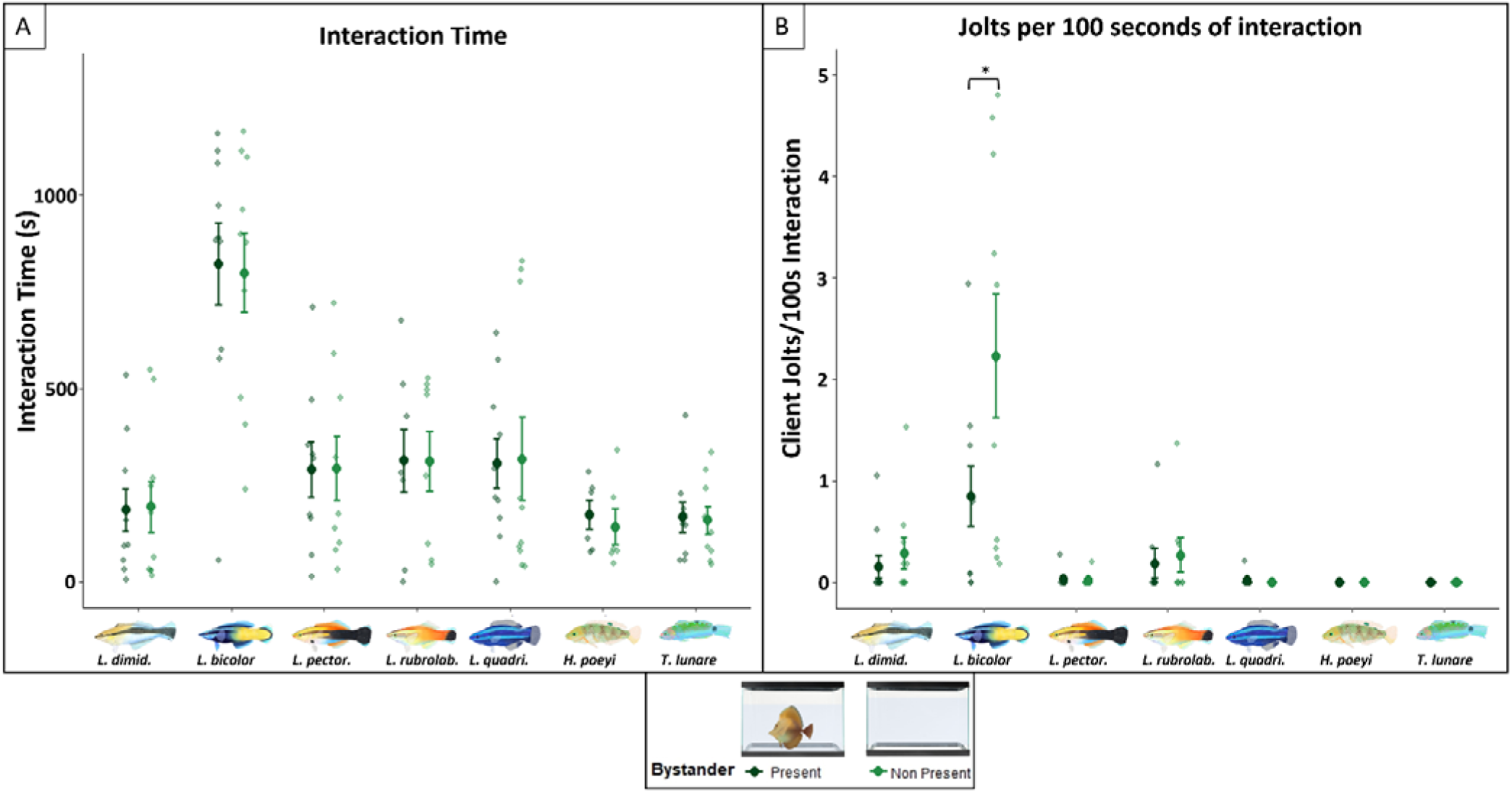
Effect of bystander presence (Present vs. Non present) on interaction time and client jolts across cleaner fish species. Panels show (A) interaction time (s) and (B) client jolts per 100 s of interaction. Small points represent individual mean values (one per cleaner individual), large points show the species mean, with error bars indicating ± SE. Colour denotes bystander condition (dark green = Present; light green = Non present). Asterisks indicate significant differences between bystander conditions within a species (paired Wilcoxon signed-rank test; * p < 0.05).

## Discussion

We investigated interspecific variation in cheating behaviour, measured via client jolts as a behavioural proxy, among four dedicated and three non-dedicated cleaner species, using two behavioural assays that measured responses to different client types and the presence of a bystander. Our results reveal that, under standardised experimental conditions, cleaner species responded differently to the same social stimuli, highlighting distinct interspecific variation in cleaning behaviour, with dedicated cleaners differing strongly among species in interaction time and client jolts, whereas non-dedicated cleaners showed broadly similar patterns characterized by lower total interaction time and rare client jolts. Dedicated cleaners also showed client-type differences across both measures, indicating that exploitation and investment can vary with partner identity. Overall, bystander effects were weak and inconsistent across species; however, *L. bicolor* showed a significant reduction in client jolts when a bystander was present, indicating that audience sensitivity may be expressed in a species-specific manner.

Our multispecies experimental framework suggests that cleaners are not functionally interchangeable, thus one species cannot be assumed to provide the same service profile as another, because species differ consistently in how they couple service investment (interaction time) with service quality (jolt occurrence and, when present, jolt rates). The markedly greater behavioural variance among dedicated cleaners is consistent with the idea that strong reliance on cleaning can impose persistent selection on the decision rules that balance service investment and exploitation (Bshary & Oliveira, 2015), and that even closely related dedicated cleaners can diverge substantially in how they deliver cleaning services (Côté & Brandl 2021). In reef systems, opportunities to interact with clients may be abundant yet contested in time and space, and diversification in service delivery could reduce direct overlap among sympatric cleaners, through variation in interaction investment, client-type targeting, or the balance between cooperative service and potential exploitation, thereby facilitating coexistence and stabilizing the mutualism at the community level (Côté & Brandl, 2021; Côté & Mills, 2020). Importantly, this variation also implies that clients are not passive recipients of service, higher jolt occurrence/rates indicate more costly interactions that can erode client net benefits and ultimately impact fitness (Bshary, 2002). Clients can therefore enforce partner control by leaving, switching, or reacting aggressively after jolts, generating feedback that may favour different investment-exploitation pattern across cleaner species and be especially consequential for dedicated cleaners, consistent with their pronounced behavioural variance (Grutter & Bshary, 2003; Bshary, 2002). By contrast, non-dedicated cleaners exhibited comparatively uniform patterns, interactions were consistently short and client jolts were rare and sporadic, consistent with weaker and less sustained selection on service delivery in species that engage in cleaning opportunistically, and thus potentially reduced pressure for diversification in how service is provided (Côté & Brandl, 2021; Barbu et al., 2011). Therefore, these patterns suggest that reliance on cleaning may broaden the range of viable service phenotypes in dedicated cleaners, while non-dedicated cleaners remain constrained to shorter and low-jolt interactions, potentially experiencing weaker selection for divergence in service delivery.

At finer resolution, these differences play out differently across client types, with species diverging in both interaction time and client jolts. *L. bicolor*, for example, showed the highest jolt expression across client types, suggesting a strategy aimed at maximising short-term rewards, whereas *L. rubrolabiatus*, consistent with its reputation for high cooperativeness in the wild, kept jolt levels consistently low across client contexts (Mills & Côté, 2010; Côté & Mills, 2020). Notably, across all cleaners, jolt rates were modest in absolute terms compared with those typically reported for wild cleaning interactions (Mills & Côté, 2010; Barbu et al., 2011; Bshary, 2002; Côté & Mills, 2020), suggesting that captive conditions can yield behavioural outcomes that differ from those expressed under natural ecological constraints. This shift was also evident in *L. bicolor*, which showed the longest interaction times among cleaners; this contrasts with field reports of relatively brief interactions in the wild (Côté & Brandl, 2021) and suggests that captivity may relax constraints shaping cleaning investment (i.e., how much time cleaners devote to clients). In line with this, predator interactions in our study showed higher jolt expression than other client contexts, most notably in *L. bicolor*, and to a lesser extent in *L. dimidiatus* and *L. pectoralis*, in contrast to field observations where predator interactions typically show very low jolt expression (Bshary, 2002). This divergence may reflect how the balance of costs and constraints shifts under captive conditions, allowing client context to reshape service outcomes even in species that are usually considered strongly regulated by partner control in the wild. Furthermore, across cleaners, interactions with the resident client were characterised by lower investment and minimal jolt expression, and even *L. bicolor* reduced both interaction time and client jolts in this context. This pattern may reflect the resident’s small body size (∼4 cm), making it a lower-value partner, consistent with field evidence that *L. bicolor* preferentially interacts with larger client species (Côté & Brandl, 2021). More broadly, it mirrors a payoff-dependent shift documented in *L. dimidiatus*, where cleaners appear more cooperative in encounters with small resident clients while increasing exploitation toward larger, more rewarding clients (Bshary, 2002). Overall, these results indicate that service allocation is client-dependent and may be reshaped in captivity, while also revealing species-specific variation in how cleaners adjust interaction time and jolt expression across standardised client contexts.

Regarding non-dedicated cleaners, our findings indicate that they engage in cleaning behaviours less frequently and exhibit lower instances of cheating compared to dedicated cleaners. Within this generally low-jolt profile, however, species still differed in the extent to which they appeared to rely on cleaning interactions. *L. quadrilineatus* stood out by showing the longest interaction times among non-dedicated cleaners, consistent with a more structured and ecologically meaningful cleaning strategy than the other non-dedicated species (Barbu et al., 2011). This aligns with field observations in sympatric populations, where *L. quadrilineatus*, despite being classified as non-dedicated, is among the most cleaning-reliant non-dedicated species: it cleans frequently, regularly interacts with predatory clients, and uses tactile stimulation, traits considered markers of more sophisticated strategic behaviour in cleaning mutualisms (Barbu et al., 2011). By contrast, *T. lunare* and *H. poeyi* interacted only briefly with clients in our assays, consistent with a more opportunistic, low-investment service profile (Roede, 1972; Barbu et al., 2011). Furthermore, echoing the divergence from wild patterns observed in dedicated cleaners, both *T. lunare* and *L. quadrilineatus* elicit higher client jolt expression during cleaning interactions in the wild (Barbu et al., 2011), whereas in our captive assays jolts were largely absent, reinforcing the idea that the experimental context may dampen or reshape key interaction outcomes relative to those reported under natural ecological constraints. Overall, variation among non-dedicated cleaners appears to be expressed primarily through differences in how strongly species invest in cleaning interactions, with *L. quadrilineatus* showing signature of higher cleaning reliance and more structured service delivery, while *T. lunare* and *H. poeyi* remain closer to an opportunistic, low-investment profile, an outcome that may be especially sensitive to the altered costs and constraints of captivity.

Across species, the presence of a bystander produced only modest shifts in cleaning outcomes, suggesting limited audience sensitivity under our experimental conditions. Bystander presence affected behaviour most clearly in *L. bicolor*, which showed a significant reduction in client jolt expression when an audience was present, indicating a context-dependent adjustment of service quality. This aligns with evidence that *L. bicolor* can flexibly modulate cooperation in relation to expected future payoffs, reducing cheating when repeated encounters with clients are more likely (Oates, Manica and Bshary, 2010). In the other species, bystander effects were weak, which may reflect the ecological context of the source populations, as high client-to-cleaner ratios, where clients are abundant and competition among cleaners is low, may reduce selection for audience-dependent behavioural adjustment (Triki et al., 2018; 2019a; 2019b). By contrast, socially complex environments, where cleaners compete for clients and maintain longer-term relationships, are more likely to favour the evolution and expression of audience sensitivity (Triki et al., 2018; 2019a; 2019b). Nevertheless, the directional reduction in jolts observed in *L. dimidiatus* and *L. rubrolabiatus* suggests that, even if the effect was not statistically supported, a larger sample size could increase power to detect such subtle behavioural adjustments. Among non-dedicated cleaners, no evidence of bystander sensitivity was detected, which could be attributed to a lack of selective pressure to monitor audiences in species that clean sporadically or primarily during early life stages (Barbu et al., 2011; Roede, 1972). Therefore, these results suggest that *L. bicolor* can express socially sophisticated, audience-related adjustment of service quality under standardised experimental conditions, and highlight that audience sensitivity may be species-specific and shaped by ecological context.

In the wild, however, cleaners operate within familiar specific local assemblages and interact repeatedly with recognizable client individuals; such long-term familiarity, together with multiple simultaneous or sequential bystanders and queueing at cleaning stations, which can shape incentives for cooperation and reputation management in ways that may not be fully captured in captivity (Bshary & Grutter, 2002b; Côté & Brandl, 2021; Grutter, 1996). While our experimental design provides control and standardization; several features of natural systems are unavoidably simplified. Additionally, species may vary in their sensitivity to captive conditions, and some caution when comparing behaviours across taxa is warranted. Lastly, a caveat of this study is that we did not directly quantify parasite loads on client fish, a factor known to influence both the expression and detectability of cheating. However, all clients were deparasitized and maintained under the same housing conditions for an extended period, and the availability of parasites, dead skin, and damaged tissue was likely similar across individuals. Therefore, while client jolts are commonly used as a proxy for cheating behaviour, they should still be interpreted with some caution, as they may also arise from non-parasitic interactions such as exploratory biting or unsolicited contact.

Taken together, our findings suggest that cheating in cleaner wrasses is shaped by a complex interplay of species-specific responses, social context, and ecological reliance on mutualism. We found that dedicated cleaners exhibited greater behavioural variability and client-specific responses, suggesting stronger selective pressures to maintain stable cooperative interactions. In contrast, non-dedicated species showed limited behavioural modulation, likely reflecting a reduced investment in cleaning and fewer incentives to develop socially sophisticated behaviours. Overall, our results show that cleaner wrasses do not respond uniformly to the same standardised social stimuli, highlighting uneven expression of social-context-dependent behavioural modulation across species in cleaning mutualisms. More broadly, this supports the view that behavioural diversity within mutualisms can emerge from divergent responses to shared social challenges, with important implications for the evolution and stability of cooperative interactions.

## Supporting information

Sypplementary Tables

## Acknowledgements

We thank all members of the Schunter Lab and the Behavioural Ecology and Evolution Group for the stimulating discussions and valuable feedback throughout the development of this study. We extend special thanks to Hugo Lee, who illustrated all the fish depicted in this work.

## >CRediT

Conceptualization: DR, CS, JRP, Data curation DR; Formal analysis DR; Funding acquisition JRP, CS; Investigation DR, MR, MC, BP; Methodology DR, CS, JRP; Project administration CS, JRP; Resources CS, JRP; Software DR; Supervision CS, JRP; Validation DR; Visualization DR; Writing – original draft DR, CS; Writing – review & editing DR, MR, MC, BP, JRP, CS.

## Ethics

All experimental procedures were conducted in accordance with institutional and governmental guidelines for the care and use of animals in research. Ethical approval for this study was granted by the Faculdade de Ciências da Universidade de Lisboa animal welfare committee (ORBEA – Statement 4/2023) and Direção-Geral da Alimentação e Veterinária (DGAV permit 27950/25-S) in accordance with the requirements imposed by the Directive 2010/63/EU on the protection of animals used for scientific purposes. Ethical approval was also granted by the Committee on the Use of Live Animals in Teaching and Research (CULATR) of The University of Hong Kong (Protocol number: 23-258) and by the Department of Health of the Hong Kong Government (Ref. number: 23-521).

## Data Availability Statement

The data supporting the findings of this study are available in Figshare at https://figshare.com/s/ab8eb3c8c141a8dc997e

## Bibliography

1. Albert, A., & Anderson, J. A. (1984). On the existence of maximum likelihood estimates in logistic regression models. Biometrika, 71(1), 1–10. 10.2307/2336390

2. Baliga, V. B., & Law, C. J. (2016). Cleaners among wrasses: Phylogenetics and evolutionary patterns of cleaning behavior within Labridae. Molecular Phylogenetics and Evolution, 94, 424–435. 10.1016/j.ympev.2015.09.006

3. Baliga, V. B., & Mehta, R. S. (2019). Morphology, Ecology, and Biogeography of Independent Origins of Cleaning Behavior around the World. Integrative and Comparative Biology, 59(3), 625– 637. 10.1093/icb/icz030

4. Barbu, L., Guinand, C., Bergmüller, R., Alvarez, N., & Bshary, R. (2011). Cleaning wrasse species vary with respect to dependency on the mutualism and behavioural adaptations in interactions. Animal Behaviour, 82(5), 1067–1074. 10.1016/j.anbehav.2011.07.043

5. Bronstein, J. L. (1994). Our Current Understanding of Mutualism. The Quarterly Review of Biology, 69(1), 31–51. 10.1086/418432

6. Brooks, M. E., Kristensen, K., Benthem, K. J. van, Magnusson, A., Berg, C. W., Nielsen, A., Skaug, H. J., Mächler, M., & Bolker, B. M. (2017). glmmTMB Balances Speed and Flexibility Among Packages for Zero-inflated Generalized Linear Mixed Modeling. The R Journal, 9(2), 378. 10.32614/RJ-2017-066

7. Bshary, R. (2002). Biting cleaner fish use altruism to deceive image–scoring client reef fish. Proceedings of the Royal Society of London. Series B: Biological Sciences, 269(1505), 2087–2093. 1010.1098/rspb.2002.2084

8. Bshary, R., & Bronstein, J. L. (2011). A General Scheme to Predict Partner Control Mechanisms in Pairwise Cooperative Interactions Between Unrelated Individuals. Ethology, 117(4), 271–283. 10.1111/j.1439-0310.2011.01882.x

9. Bshary, R., & Grutter, A. S. (2002a). Asymmetric cheating opportunities and partner control in a cleaner fish mutualism. Animal Behaviour, 63(3), 547–555. 10.1006/anbe.2001.1937

10. Bshary, R., & Grutter, A. S. (2002b). Experimental evidence that partner choice is a driving force in the payoff distribution among cooperators or mutualists: the cleaner fish case. Ecology Letters, 5(1), 130–136. 10.1046/j.1461-0248.2002.00295.x

11. Bshary, R., & Oliveira, R. F. (2015). Cooperation in animals: toward a game theory within the framework of social competence. Current Opinion in Behavioral Sciences, 3, 31–37. 10.1016/j.cobeha.2015.01.008

12. Côté, I. M., & Brandl, S. J. (2021). Functional niches of cleanerfish species are mediated by habitat use, cleaning intensity and client selectivity. Journal of Animal Ecology, 90(12), 2834–2847. 10.1111/1365-2656.13585

13. Côté, I. M., & Mills, S. C. (2020). Degrees of honesty: cleaning by the redlip cleaner wrasse Labroides rubrolabiatus. Coral Reefs, 39(6), 1693–1701. 10.1007/s00338-020-01996-6

14. Cragg, J. G. (1971). Some statistical models for limited dependent variables with application to the demand for durable goods. Econometrica: journal of the Econometric Society, 829-844. 10.2307/1909582

15. de Abreu, M. S., Messias, J. P. M., Thörnqvist, P.-O., Winberg, S., & Soares, M. C. (2018). The variable monoaminergic outcomes of cleaner fish brains when facing different social and mutualistic contexts. PeerJ, 6, e4830. 10.7717/peerj.4830

16. Ferrière, R., Gauduchon, M., & Bronstein, J. L. (2007). Evolution and persistence of obligate mutualists and exploiters: competition for partners and evolutionary immunization. Ecology Letters, 10(2), 115–126. 10.1111/j.1461-0248.2006.01008.x

17. Friard, O., & Gamba, M. (2016). BORIS: a free, versatile open source event logging software for video/audio coding and live observations. Methods in Ecology and Evolution, 7(11), 1325–1330. 10.1111/2041-210X.12584

18. Fox, J., & Weisberg, S. (2018). An R Companion to Applied Regression_, Third edition. Sage, Thousand Oaks CA. <https://www.john-fox.ca/Companion/>.

19. Goodwin, N. L., Choong, J. J., Hwang, S., Pitts, K., Bloom, L., Islam, A., Zhang, Y. Y., Szelenyi, E. R., Tong, X., Newman, E. L., Miczek, K., Wright, H. R., McLaughlin, R. J., Norville, Z. C., Eshel, N., Heshmati, M., Nilsson, S. R. O., & Golden, S. A. (2024). Simple Behavioral Analysis (SimBA) as a platform for explainable machine learning in behavioral neuroscience. Nature Neuroscience, 27(7), 1411–1424. 10.1038/s41593-024-01649-9

20. Ghoul, M., Griffin, A. S., & West, S. A. (2014). Toward an evolutionary definition of cheating. Evolution, 68(2), 318–331. 10.1111/evo.12266

21. Grutter, A. S. (1996). Parasite removal rates by the cleaner wrasse Labroides dimidiatus. Marine Ecology Progress Series, 130, 61–70. 10.3354/meps130061

22. Grutter, A. S. (2004). Cleaner Fish Use Tactile Dancing Behavior as a Preconflict Management Strategy. Current Biology, 14(12), 1080–1083. 10.1016/j.cub.2004.05.048

23. Grutter, A. S., & Bshary, R. (2003). Cleaner wrasse prefer client mucus: support for partner control mechanisms in cleaning interactions. Proceedings of the Royal Society of London. Series B: Biological Sciences, 270(suppl_2). 10.1098/rsbl.2003.0077

24. Hartig, F. (2024). DHARMa: Residual diagnostics for hierarchical (multi-level / mixed) regression models (R package version 0.4.7). https://CRAN.R-project.org/package=DHARMa

25. Irwin, R. E., Brody, A. K., & Waser, N. M. (2001). The impact of floral larceny on individuals, populations, and communities. Oecologia, 129(2), 161–168. 10.1007/s004420100739

26. Burkle, L. A., Irwin, R. E., & Newman, D. A. (2007). Predicting the effects of nectar robbing on plant reproduction: implications of pollen limitation and plant mating system. American Journal of Botany, 94(12), 1935–1943. 10.3732/ajb.94.12.1935

27. Kosmidis I. (2021). brglm: Bias Reduction in Binary-Response Generalized Linear Models. R package version 0.7.2, https://cran.r-project.org/package=brglm

28. Lenth, R. (2024). emmeans: Estimated marginal means, aka least-squares means (R package version 1.10.6). https://CRAN.R-project.org/package=emmeans

29. Levene, H. (1960). Robust tests for equality of variances. In Contributions to Probability and Statistics: Essays in Honor of Harold Hotelling (I. Olkin, Ed.), pp. 278–292. Stanford University Press.

30. Lin, H. et al. (2024) ‘Interspecific competition prevents the proliferation of social cheaters in an unstructured environment’, The ISME Journal, 18(1). 10.1093/ismejo/wrad038

31. Mills, S. C., & Côté, I. M. (2010). Crime and punishment in a roaming cleanerfish. Proceedings of the Royal Society B: Biological Sciences, 277(1700), 3617–3622. 10.1098/rspb.2010.0941

32. Nath, T., Mathis, A., Chen, A. C., Patel, A., Bethge, M., & Mathis, M. W. (2019). Using DeepLabCut for 3D markerless pose estimation across species and behaviors. Nature Protocols, 14(7), 2152–2176. 10.1038/s41596-019-0176-0

33. Oates, J., Manica, A., & Bshary, R. (2010). The shadow of the future affects cooperation in a cleaner fish. Current Biology, 20(11), R472–R473. 10.1016/j.cub.2010.04.022

34. Paula, J. R., Baptista, M., Carvalho, F., Repolho, T., Bshary, R., & Rosa, R. (2019). The past, present and future of cleaner fish cognitive performance as a function of CO 2 levels. Biology Letters, 15(12), 20190618. 10.1098/rsbl.2019.0618

35. Pellmyr, O., & Leebens Mack, J. (2000). Reversal of Mutualism as a Mechanism for Adaptive Radiation in Yucca Moths. The American Naturalist, 156(S4), S62–S76. 10.1086/303416

36. Pinto, A., Oates, J., Grutter, A., & Bshary, R. (2011). Cleaner Wrasses Labroides dimidiatus Are More Cooperative in the Presence of an Audience. Current Biology, 21(13), 1140–1144. 10.1016/j.cub.2011.05.021

37. Poulin, R. and Grutter, A.S. (1996) ‘Cleaning Symbioses: Proximate and Adaptive Explanations’, BioScience, 46(7), pp. 512–517. Available at: 10.2307/1312929.

38. Ramírez-Calero, S., Paula, J. R., Otjacques, E., Ravasi, T., Rosa, R., & Schunter, C. (2023). Neuromolecular responses in disrupted mutualistic cleaning interactions under future environmental conditions. BMC Biology, 21(1), 258. 10.1186/s12915-023-01761-5

39. Ramírez-Calero, S., Paula, J. R., Otjacques, E., Rosa, R., Ravasi, T., & Schunter, C. (2022). Neuro-molecular characterization of fish cleaning interactions. Scientific Reports, 12(1). 10.1038/s41598-022-12363-6

40. Roede, M. J. (1972). Color as related to Size, Sex, and Behavior in seven Caribbean Labrid Fish Species (genera Thalassoma, Halichoeres and Hemipteronotus). In Studies on the Fauna of Curaçao and other Caribbean Islands (1st ed., Vol. 42, pp. 1–264). The Hague: Martinus Nijhoff. https://repository.naturalis.nl/pub/506053

41. Ripley, B., Venables, B., Bates, D. M., Hornik, K., Gebhardt, A., Firth, D., & Ripley, M. B. (2013). Package ‘mass’. Cran r, 538(113-120), 822.

42. R Core Team (2023). _R: A Language and Environment for Statistical Computing_. R Foundation for Statistical Computing, Vienna, Austria. https://www.R-project.org/

43. Sazima, I., Moura, R. L., & Gasparini, J. L. (1998). The wrasse Halichoeres cyanocephalus (labridae) as a specialized cleaner fish. in bulletin of marine science (Vol. 63, Issue 3).

44. Triki, Z., Levorato, E., McNeely, W., Marshall, J., & Bshary, R. (2019a). Population densities predict forebrain size variation in the cleaner fish Labroides dimidiatus. Proceedings of the Royal Society B: Biological Sciences, 286(1915), 20192108. 10.1098/rspb.2019.2108

45. Triki, Z., Wismer, S., Levorato, E., & Bshary, R. (2018). A decrease in the abundance and strategic sophistication of cleaner fish after environmental perturbations. Global Change Biology, 24(1), 481–489. 10.1111/gcb.13943

46. Triki, Z., Wismer, S., Rey, O., Ann Binning, S., Levorato, E., & Bshary, R. (2019b). Biological market effects predict cleaner fish strategic sophistication. Behavioral Ecology, 30(6), 1548–1557. 10.1093/beheco/arz111

47. Vaughan, D. B., Grutter, A. S., Costello, M. J., & Hutson, K. S. (2017). Cleaner fishes and shrimp diversity and a re evaluation of cleaning symbioses. Fish and Fisheries, 18(4), 698–716. 10.1111/faf.12198

48. Wilcoxon, F. (1945). Individual comparisons by ranking methods. Biometrics bulletin, 1(6), 80–83. 10.2307/3001968

49. Wechsler, D., & Bascompte, J. (2022). Cheating in Mutualisms Promotes Diversity and Complexity. The American Naturalist, 199(3), 393–405. 10.1086/717865

